# Epizootic of highly pathogenic H7N3 Avian Influenza in an ecologic reserve in Mexico

**DOI:** 10.1101/2020.03.05.978502

**Authors:** Roberto Navarro-López, Mario Solís-Hernández, Miguel A. Márquez-Ruiz, Abel Rosas-Téllez, Carlos A. Guichard-Romero, Gerardo de J. Cartas-Heredia, Romeo Morales-Espinosa, Héctor E. Valdez-Gómez, Claudio L. Afonso

**Affiliations:** Mexico-USA Commission for the prevention of F & M Disease and other Exotic Diseases in Animals. DGSA-SENASICA. Av. Cuauhtémoc No. 1230 piso 9, Col. Santa Cruz Atoyac. C.P. 03310 Ciudad de México; Faculty of Veterinary Medicine. UNAM. Ciudad Universitaria, C.P. 04510. Ciudad de México; Regional Zoo “Miguel Álvarez del Toro” SEMAHN. Calzada Cerro Hueco S/N, El Zapotal C.P. 29094. Tuxtla Gutiérrez, Chiapas. México; Southeast Poultry Research Laboratory, ARS-USDA, 934 College Station Road Athens, GA 30605 USA

## Abstract

This report includes a 2015 epizootic of highly pathogenic H7N3 avian influenza virus among captive and wild birds at “El Zapotal” ecologic reserve, located in the state of Chiapas, Mexico. Epidemiological control measures were implemented to prevent virus dissemination. The infection with the highly pathogenic H7N3 virus was detected predominantly among Plain Chachalaca (*Ortalis vetula*), with occasional detections in a White-fronted Parrot (*Amazona albifrons*) and a single Clay-colored Thrush (*Turdus grayi*). Here, we describe the characteristics of the outbreak environment, the surveillance strategy, the biosecurity measures, and the evaluation of the site, including external farms. These actions, timely implemented by the veterinary authorities, helped to contain the outbreak beyond the ecologic reserve. This contingency showed the importance of developing a more complete analysis of the existing risks and the challenges to implement minimal biosecurity measures in these facilities.

## Introduction

Type A influenza viruses (AIVs) typically circulate in migratory and non-migratory wild aquatic waterfowl, gulls, and shorebirds [1]. Geographic barriers have limited intercontinental exchange of AIVs resulting in continental lineages of viral diversity by regions [2, 3, 4]. They are further divided into subtypes based on the antigenic characterization of the virus’ surface glycoproteins, hemagglutinin (HA) and neuraminidase (NA). In wild birds, 16 HA and 9 NA subtypes have been identified in many different combinations which have been isolated from a total of more than 125 bird species that belong to 26 different families [1, 2, 5, 6]. AIVs maintained in wild birds periodically spill-over into domestic birds, as wild and domestic mammals, and have been postulated to contribute to pandemic human influenza viruses [7, 8, 9].

Based on pathogenicity in young, immunologically naïve chickens (*Gallus gallus domesticus*) [10], AIVs are classified as low pathogenicity (LP) or high pathogenicity (HP). LP AIVs can become established in domestic poultry with the potential to become zoonotic. LP AIVs of the H5 and H7 subtypes may develop HP in domestic poultry through the evolution of a polybasic sequence at the HA cleavage site. Domestic birds infected with HP AIVs often-present high morbidity and mortality due to systemic infection and generalized congestive-hemorrhagic condition of diverse intensity. Morbidity and mortality of wild bird infected by HP AIVs may be more variable. HP AIV can be epitheliotropic, endotheliotropic, neurotropic or pantotropic depending on host susceptibility [11, 12]. In domestic chickens, significant lesions that are consistently associated with HP AIV infections include necrosis and inflammation of organs such as heart, brain, spleen, intestinal tract, lungs, and skeletal muscles [11].

There is limited information about HPAI outbreaks in wild birds, but HPAI infections have shown a range of syndromes that can go from asymptomatic to severe symptomatic conditions. These range from respiratory and/or digestive tract infections to critical lesions in other organs including the heart, brain and pancreas [11]. The reporting of this is type of infection in wild birds has increased during the last 15 years [13, 14, 15].

In Mexico, during an epizootic in poultry, a continued transmission of HPAI H7N3 virus was found to occur in *Quiscalus mexicanus* (Fam. Icteridae, Ord. Passeriformes), for at least six weeks. The clinical condition of the *Q. mexicanus* was, prostration, ruffle feathers, diarrhea, anorexia and sudden death [14]. Similar conditions have been reported in the *Corvus macrorhynchos* and in *Pica pica* infected with a HPAI H5N1 virus [11]. In both reports, the lesions described included severe damage of the brain, heart and pancreas and the isolation of the H5N1 and H7N3 viruses from trachea, intestine and heart.

Most of the HP AIV identified in wild birds have been detected in typical reservoir species [7, 16, 17], although a diversity of additional species has also been reported to be affected; such is the case of passerine [18,19] and psittacine birds [20]. The role of passerines as virus reservoirs in nature has been gaining terrain, facilitating viral spread between wild and domestic hosts [20, 21].

This study describes the initial evidences of mortality in a wild population of Plain chachalacas (*Ortalis vetula*), following an outbreak in a natural reserve in Mexico. We also identified the presence of the HPAIV H7N3 in White-fronted Parrot (*Amazona albifrons*), and Clay-colored Thrush (*Turdus grayi*), which were reported for the first time in an ecologic reserve where a regional fauna zoo is present. We also highlighted the procedures implemented by the veterinary authorities to achieve its control [23, 25].

## Material and Methods

### Study Area

The natural reserve “El Zapotal” located in the SE portion of Tuxtla Gutiérrez capital of the State of Chiapas, Mexico, (16° 43’ N / 93° 06’ W), comprises 192 hectares of Semi-evergreen forest, Low deciduous woodland and Acacia shrub [26]. The climate was defined as warm, humid and rainy weather (Awo(w)(i)), with a total annual precipitation of 948.2 mm and average temperature of 24.7 °C [27]. The decree of protection was established as a preventive measure against the intense urban pressure. The Regional Zoo Miguel Álvarez del Toro (ZooMAT), is located within this natural reserve (Fig. 1). There is also a profusely free-ranging wild animal population, including feral fauna interacting year-round [23, 26, 28].

**Figure 1.**
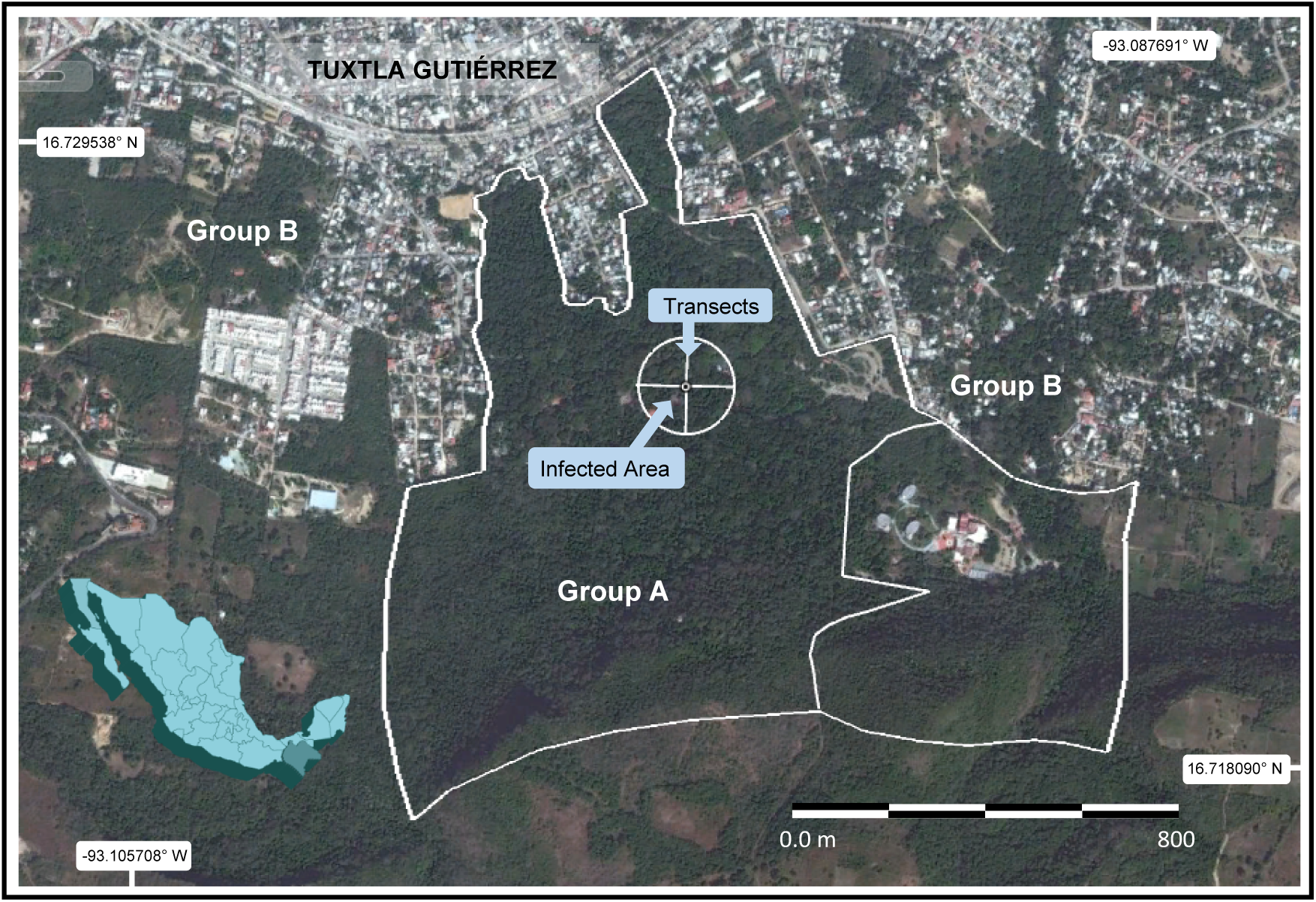
Polygon of the ecological reserve El Zapotal, including working groups zones.

### Avifauna and target species

El Zapotal, is rich in avifauna with 364 species dominated by Passeriformes. The seasonal behavior shows that 68% are resident breeders, 32% are winter visitors while 34% are considered to be rare [33]. The target species was the Plain chachalaca, whose distribution ranges from South Texas, the Gulf of Mexico, the Yucatan peninsula to North Central America [34, 36]. Within the reserve, the abundance of resources such as fruits, seeds, leaves and insects, comprises the main items present in the diet of the chachalaca [26]. This condition among other factors maintain a relatively high density of individuals [34, 35].

The Zoo, where “focal” or infected zone was designated, is surrounded by aviaries mainly populated by Psittacine birds. Some artificial ponds are also present, harboring resident aquatic birds, as well as an important diversity of songbirds [33].

### Method of Study

In order to address the epizooty and the procedures implemented, the surveillance staff was divided in three groups (A, B and C) following biosecurity protocols [22] to prevent contamination (Figure 1). Group A was responsible for the activities within the reserve consisting of: Attention to AIV suspect infected cases, delimitation of the infected area, stamping out of the birds, postmortem examination, as well as blood and organ extractions. Other activities included cleanliness and disinfection of the facilities, risk evaluation, risk control, in addition to sampling of exhibited birds via tracheal and cloacal swabbing every 21 days during three months. Group B was responsible for the communication protocols within the urban and sub-urban (perifocal) zone around El Zapotal (Fig. 2). Group C was in charge of serologic and virologic surveillance of commercial broiler farms located within 10 km perimeter around El Zapotal (Fig. 2). This group carried out a minimal biosecurity evaluation of the farms, in order to avoid cross-contamination of poultry. During this contingency, the access of official veterinarians to commercial farms was forbidden.

**Figure 2.**
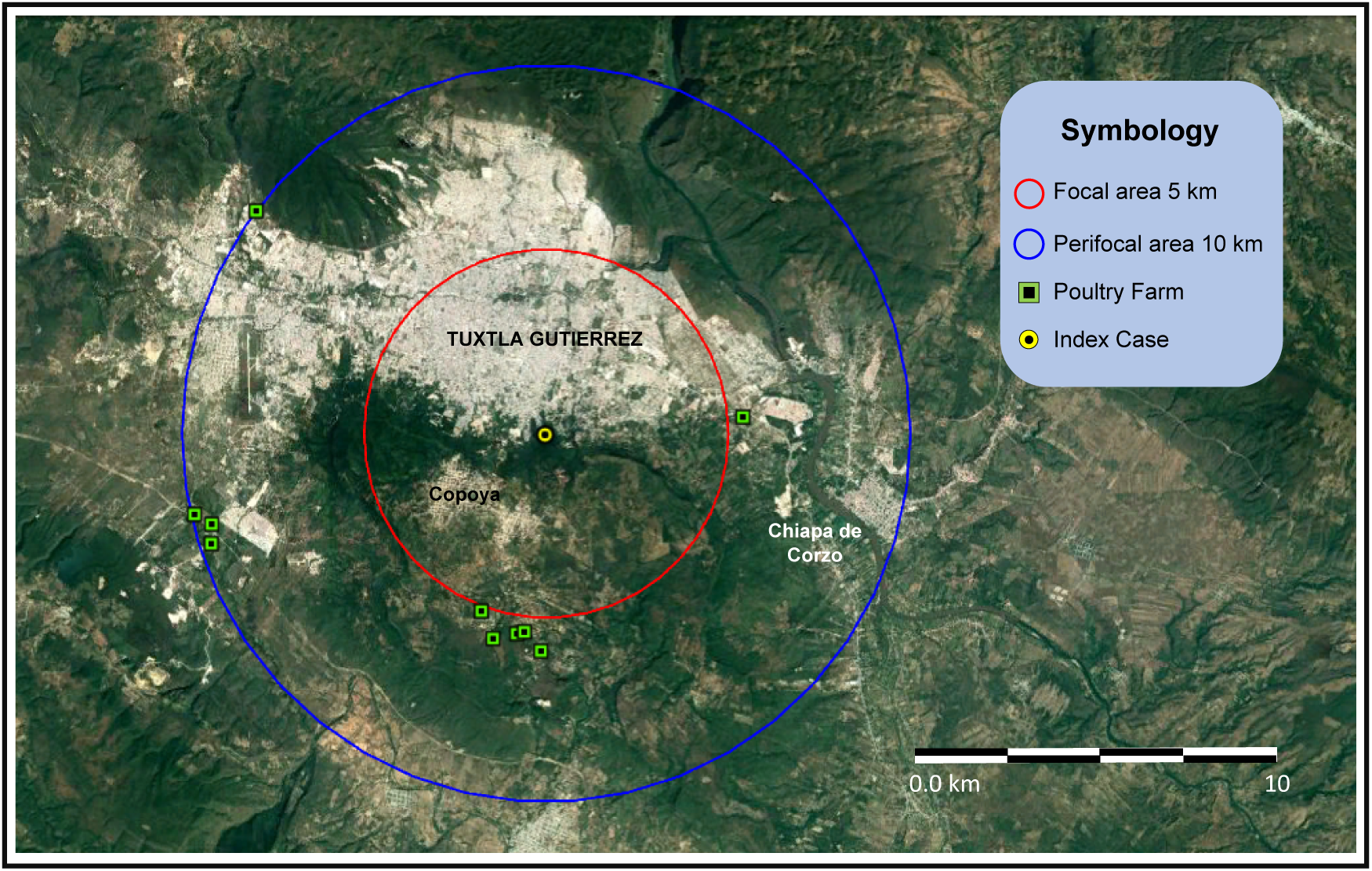
Focal area and Perifocal area around El Zapotal.

Virologic surveillance was carried out by sampling daily mortality in poultry. Tracheal and cloacal swabbing performed by official veterinary were obtained from carcasses placed in containers outside the farms. The serologic surveillance was carried out by non-official veterinarians accredited by the authority.

The infected area was delimited considering the index case (IC) as the center of a circle of 100 m radius. All chachalacas and birds showing symptomatology where shot with two air rifles type PCP Cal. 5.5. Additionally, two transects of two hundred meters each, were implemented toward the four cardinal points, by placing the IC at its intersection. Along these transects active vigilance on wild birds took place (Fig. 1). These actions were endorsed by the Articles 8, 11, and 160 of the General Law of Animal Health (24).

We calculated the minimal number of samples by implementing the Cannon & Roe formula [31]. Population density for *O. vetula* were assumed considering previous censuses [35] with a prevalence of 3%. A total number of commercial birds within the perifocal zone was applied considering a prevalence of 10% for intra-farms, with a level of confidence of 97% for both cases [29].

Protocols such as Guidelines to the Use of Wild Bird in Research were considered [30]. Poultry flock surveillance strategies, clinical examination, serologic and virologic sampling were carried out as described in the Terrestrial Animal Sanitary Code of OIE [22, 32, 42]. All field activities were implemented according to the Manual of Procedures for the prevention and eradication of HPAI virus [22].

### Phylogenetic analyzes

To examine the relationship between the HPAI H7N3 from ZooMAT, 16 full-length HA nucleotide sequences were aligned (isolated from ZooMAT, Puebla, Oaxaca, Jalisco and two additional low pathogenicity viruses). Next, a maximum likelihood tree was constructed using the Tamura-Nei model. Initial tree (s) for the heuristic search were obtained by using the Maximum Composite Likelihood (MCL) approach, and then selecting the topology with superior log likelihood value. A discrete Gamma distribution was used to model evolutionary rate differences between sites. The tree was drawn to scale with branch lengths measured in the number of substitutions per site. Evolutionary analyzes were conducted in MEGA7.

### Diagnostics tests

The samples were processed according to the Catalogue of Procedures and Services of the Laboratory Network of the DGSA [38, 39]. Samples were subjected to virus isolation in embryonated chicken eggs and reverse transcription polymerase chain reaction in real time (rRT-PCR). Virus subtype and pathogenicity were assessed by hemagglutination inhibition, sequencing, and intravenous pathogenicity index (IVPI), OIE’s method adopted.

For the determination of the nucleotides at that portion of the HA gene coding for the cleavage site region of the haemagglutinin has been used the Applied Biosystems^®^ 3130 Genetic Analyzer. The completed haemagglutinin sequence was performed by using the conserved AIV primers to amplify a product that was sequenced by the Illumina Nexgen MiniSeq™ System.

### Databases

The informatics platform used for the collection and analysis of the data was obtained from the Mexican National System of Information for Exotic and Emergent Diseases –SINEXE-(39). The number and origin of people visiting the ZooMAT along this study was taken from to the zoo’s own data.

## Results

The epidemiologic analysis showed that the IC, was the primary case (PC) detected on April 20 2015, in two *Ortalis vetula* by RT-PCR technique; both birds had died on April 16. On April 29, viral isolation and identification confirmed the existence of AIV H7N3 subtype. On April 30, this virus was sequenced and on May 4, the results of Intravenous Pathogenicity Index (IVPI) exposed 2.91-2.99 (100% mortality) indicating a HPAI H7N3 virus. The clinical signs of inoculated chicken exposed to the disease for the IVPI were hyperacute. These included ruffled feathers, respiratory distress, nasal discharge, conjunctivitis, torticollis, prostration and sudden death. The macroscopic lesions observed corresponding to petechial hemorrhages in the comb, tarsi and barbels, lung congestion, hemorrhagic tracheitis, proventricular hemorrhage, and hemorrhages in the muscles of the breast.

Group A collected 1,853 virologic samples for diagnosis. Among them 511 were cloacal and tracheal swabs, 198 organs, 1,135 dragging swabs, and 9 sera (Table1). From these samples, 64 birds were positive to virus isolation. Positive samples belonged to three species grouped in three taxonomic families (Table 2).

**Table 1.** Number of samples collected, positive to Highly Pathogenic Avian Influenza H7N3, per group.

| Team | Number of Samples | HPAI H7N3 |
| --- | --- | --- |
| Group A | 1,853 | 64 |
| Group B | 702 | 0 |
| Group C | 2,523 | 0 |
| **Total** | 5,078 | 64 |

**Table 2.**
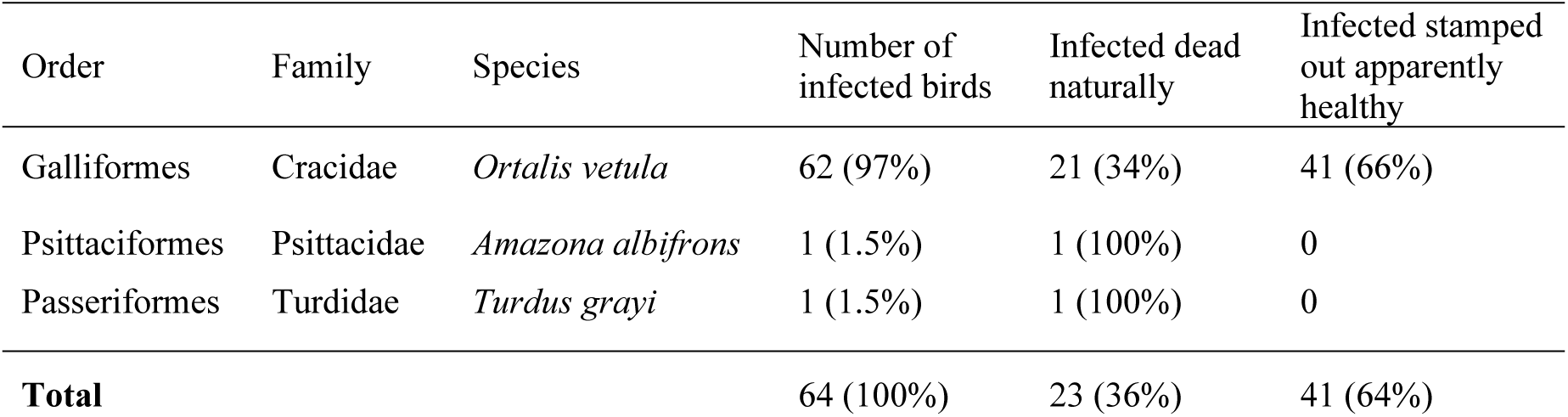
Taxonomic description and proportions of infected wild birds.

The minimal sample size calculated suggested 145 individuals with Finite Population Correction. The infected area had 3.14 ha where a total of 193 birds were sacrificed (depopulated). 174 *Ortalis vetula*, 6 *Turdus grayi*, 5 *Molothrus aeneus*, 4 *Egretta thula*, 2 *Coragyps atratus* 1 *Quiscalus mexicanus* and 1 *Amazona albifrons*. One hundred percent of the birds were sampled for AIV, 64 were positive for H7N3: 62 *Ortalis vetula*, 1 *Turdus grayi* and 1 *Amazona albifrons* (Table 2).

Group B. The surveillance teams detected 3,141 locations with backyard birds, predominately represented by *Gallus gallus domesticus*, although, pigeons, ducks, pheasants, peacocks, turkeys, parrots and songbirds were also identified. Through interviews and visual inspection, it was determined that *O. vetula* was able to come in contact with poultry and backyard birds in the neighboring sites that surround the “El Zapotal” natural reserve. During the research, we received 53 notifications of ill birds by different causes. A total of 702 samples were collected: 45 organs, 657 tracheal and cloacal swabs, however none of the samples collected had the A H7N3 virus (Table 1).

Group C. This group carried out biosecurity evaluation of the farms. Around the ecologic reserve, we identified 11 commercial farms (Fig. 2), 8 broilers and 3 rearing farm operations. Under the dead-bird sampling system, 1,743 tracheal and cloacal swabs were obtained in 83 visits and the accredited veterinarians also collected 780 blood samples (Table 1). No evidence of H7N3 virus presence and/or exposure was detected. To verify compliance with minimal biosecurity practices, 29 farms were audited, 10 in the perifocal area (Fig 2) and 19 in the area where they wanted to improve the biosecurity conditions. Among these farms, 21 were considered to have in place appropriate biosecurity measures, whereas 8 were required to improve their preventive infrastructure.

64% of the cases were found by controlled depopulation and 36% were detected through passive surveillance with carcasses. The ratio of infected males *vs* females in *O. vetula* was 3 to 1. The epizootic lasted 6 weeks from April 17 (IC) to May 23, which corresponds to epidemiologic weeks 16 to 21. The sampling of *O. vetula* started on week 18 and finished on week 26. (Fig. 3).

**Figure 3.**
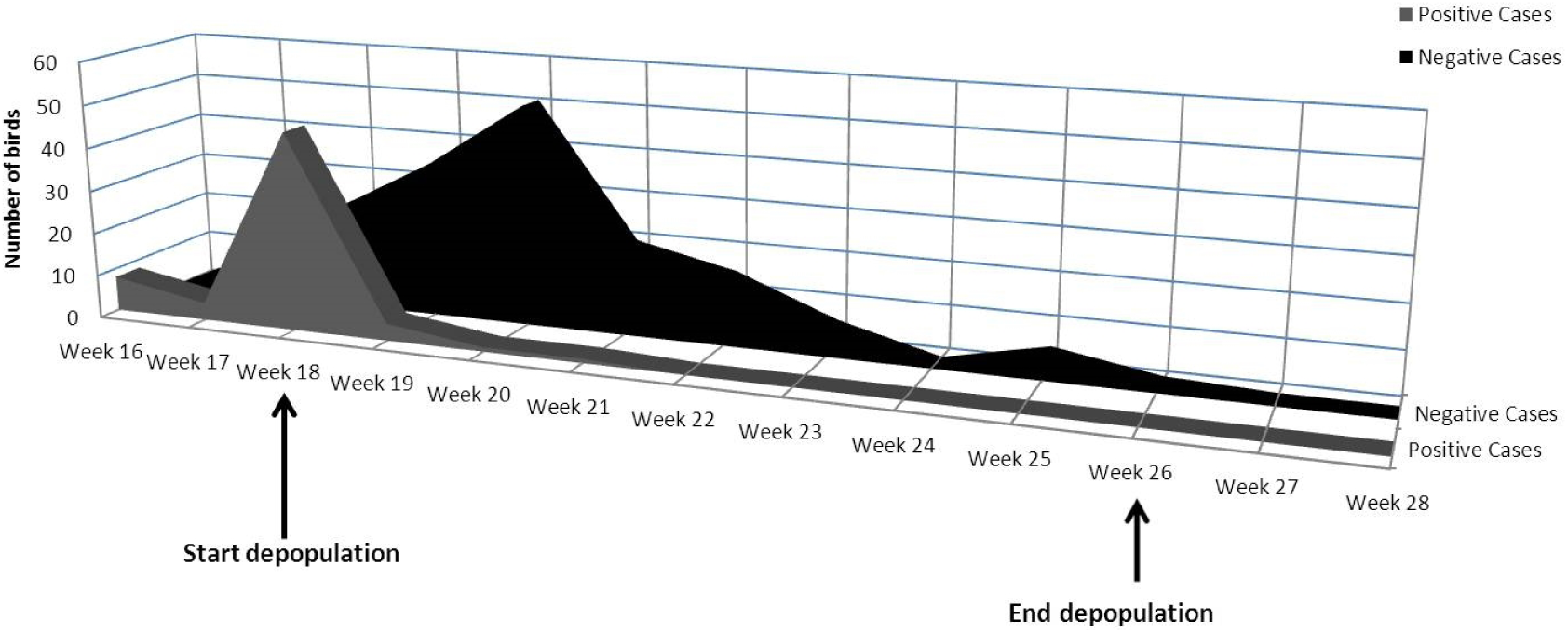
Chronology of the outbreak of Avian Influenza H7N3 HP in the ecological reserve El Zapotal.

### Clinical signs and post-mortem findings

Most of the infected birds with the HPAI H7N3 virus died rapidly in less than 24 hours (Fig. 4B). Clinically ill birds showed signs of depression, ruffled feathers, nasal discharge, anorexia, dehydration, yellow diarrhea, fever (43°C), most likely due to severe systemic infection (Fig. 4A). During post-mortem examination a generalized congestive-hemorrhagic condition was observed with the following findings: air sacs opacity, heart necrosis, necrotic foci in liver, tracheal congestion and hydropericardium, gastroenteritis, hemorrhagic enteritis, and splenomegaly, presence of bloody fluids in the coelomic cavity (Fig. 4 C-D).

**Figure 4.**
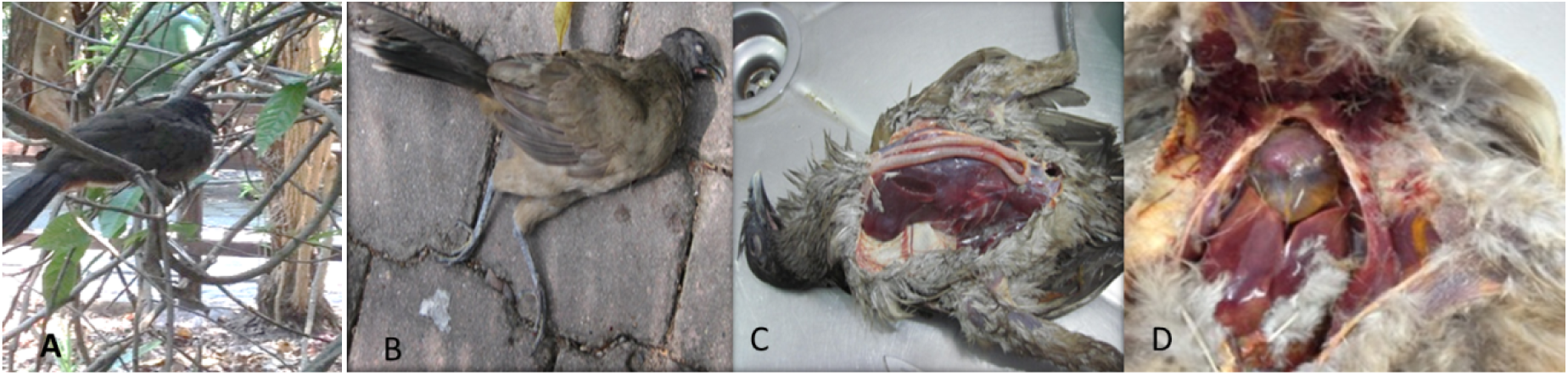
*Ortalis vetula* affected with HPAI H7N3 virus.

### Sources of the outbreak

According to the SINEXE, there were no receptions on the database of dead birds in the ZooMat since April 1-15, suggesting that the virus was probably not present on the premises before the IC was detected. Based on the list of visitors 27,294 people attended the zoo during April 1-15, 80% originated from Chiapas and the rest from other Mexican states.

### Control of the outbreak

ZooMAT implemented a stringent monitoring and control program based on clinical and virological surveillance, previously verified by the veterinarian authorities according to OIE as a consequence of a Newcastle Disease outbreak [40, 41]. The actions implemented at the zoo were the following: Temporary closure and depopulation of the *O. vetula* in the focal area, in order to curtail the transmission chain. In addition, since 2008 a program of surveillance for Avian Influenza and Newcastle viruses has been established using PCR molecular tests and viral isolation on birds that come to the zoo through donations, confiscation or birds that die within the zoo.

The infected area was cleaned with water and soap and disinfected with citric acid (2%, commercial name Desfan-100) using a pressure pump. Organic materials were sprayed with Desfan-100 during 14 days and then were composted in situ. At the two zoo entrances, pumps and footbaths with disinfectant for footwear were placed and vehicles were disinfected with active neutral (pH 0.006%) chloraldehyde (1%). Footbaths were also placed in the aviaries using the same disinfectant and concentration. Same actions were applied at the three points around infected area inside the zoo. Covered garbage cans were used to replace those without cover, in order to prevent feral birds and O. vetula from having contact with trash and leftover food at the two restaurants of the zoo. High-risk materials, such as organic waste from the restaurants and dead birds were destroyed in the incinerator located within the zoo. During the 90 days of active surveillance, 100% of the exhibition aviaries were sampled (1,264 swabs obtained from 500 clinically healthy birds) with negative results for both rRT-PCR and virus isolation.

The tree with the highest log likelihood (−3494.0680) is shown. The percentage of trees in which the associated taxa clustered together is shown next to the branches. The discrete Gamma distribution used to model evolutionary rate differences between 5 sites (5 categories: + G, parameter = 1.0429). The analysis involved 16 nucleotide sequences. All positions containing gaps and missing data were eliminated. There were a total of 1683 positions in the final dataset (Fig. 5).

**Figure 5.**
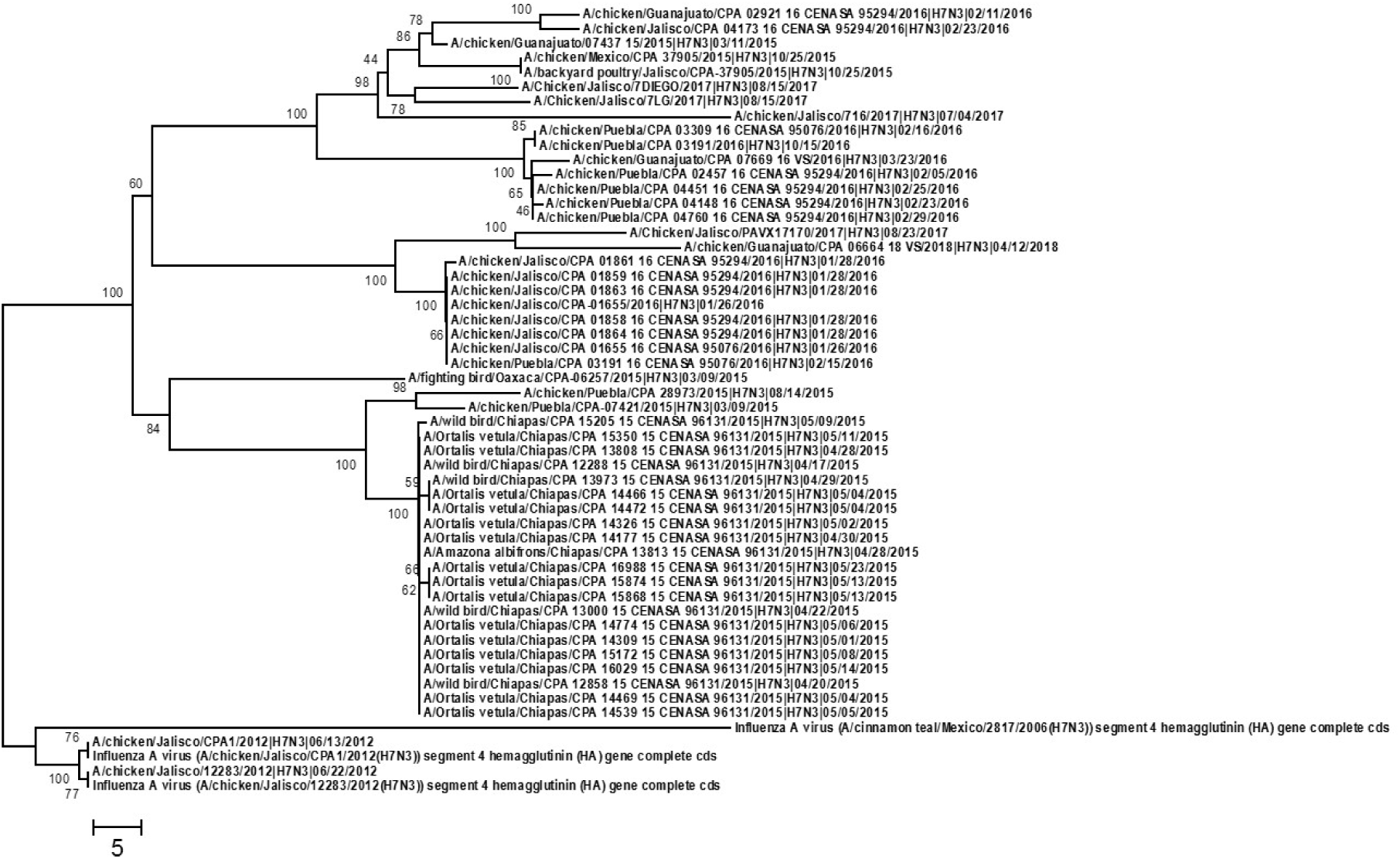
Evolutionary relationships of taxa from HPAI H7N3 isolated from *Ortalis vetula*, located in ZooMAT, Chiapas.

## Discussion

Here we reported the first case of infection of *Ortalis vetula, Turdus grayi* and *Amazona albifrons* with HPAI H7N3 virus, in the “El Zapotal” natural reserve. In addition, the implementation of established procedures was used to control poultry infections in order to protect wild bird populations. The AIV epizootic was eliminated in the natural reserve, complying with the “90 days period without any new case”, required by OIE, in order to declare free a zone of Avian Influenza [42] and did remain free for at least 105 days (week since the end of the campaign. Particularly significant is the fact that *O. vetula* came in contact with backyard birds in the neighboring sites around the natural reserve. These arguments are visually based or through interviews. The high prevalence observed in infected area, demonstrates the capacity of this species to be infected by H7N3 and to efficiently transmit the virus under field conditions.

In this particular case, the biosecurity measures undertaken to control the AIV resemble poultry farms in Mexico. A vaccination program under official control is used to prevent AIV infection and spreading, accompanied by quarantine and stamping out policies according to the requirements established by the veterinarian authorities [22, 40, 43].

Extending stamping out practices to non-endangered wild bird species within their natural environment proved effectiveness in controlling the spread of the HPAI H7N3 virus in *O. vetula*, thus preventing the potential spill over into other endangered species. That is case of Resplendent Quetzal (*Pharomachrus mocinno*); Horned Guan (*Oreophasis derbianus*), parrots and macaws (*Ara* spp.). The involvement of some of these species would have resulted in a serious animal health conflict, where public opinion plays a very important role pressure. Upon early detection, the key action implemented at the ZooMAT was the rapid depopulation of *O. vetula* in the focal area. By reducing its population in this critical spot of the zoo, it may help to reduce the transmission chain. The epizootic lasted six weeks due to the implemented actions undertaken by the official veterinary services (Figure 2). During controlled depopulation, it was discovered that many apparently healthy birds were infected. The cleaning actions, disinfection, waste handling and control, the worker’s training and the improvement of biosecurity in general, were all key actions that contributed to the contention of the disease. These actions prevented wild birds to act as “bridge birds” by infecting other animal populations.

Our data further emphasizes the delicate balance that exists at the wild bird/poultry interphase (44, 45).

The ecological consequences of this process in which highly virulent viruses develop in poultry and return to the wild are unknown.

This outbreak represents yet another case in which an AIV started as a LPAI virus in wild birds becoming highly pathogenic in poultry and then circling back as HPAI virus into wild birds. The HPAI H7N3 virus was reported for the first time in Mexico in June 22, 2012 [48], in the state of Jalisco at layer hens, in places where wild migratory birds come together from different migratory routes during wintertime [46, 47].

The sequence of the hemagglutinin cleavage site had 100% similarity with the sequence of HPAIV isolated from Jalisco in 2012. As shown in the image 1, the inclusion of 24 nucleotide described by Maurer-Stroh et. al., clearly shows a widespread anchor associated with the host 28S ribosomal RNA (rRNA). The host range expansion of a HPAIV H7N3 virus was previously detected with the presence of viruses of this serotype in poultry in Mexico.

The state of Jalisco, located in west-central Mexico, provides the largest production of table eggs in the country, followed by the state of Puebla [51]. In these states, vaccination against avian influenza H7 is authorized. From 2013 the H7N3 HP virus has extended its geographical distribution [49, 50, 52, 53, 54, 55,].

The *O. vetula* population is abundant and possible overpopulated in this reserve due to plentiful food and to predator’s absence. Considering that there must be more than 1,100 individuals (5.9 individuals/ha) [34, 35]. We believe that the AI infection in *T. grayi* and *A. albifrons*, was not progressive, and that these are spillover from the infection in *O. vetula*, since we did not find evidence of ill birds or even dead ones. However, other exhibited Cracids such as Great Curassow (*Crax rubra*) and Crested Guan (*Penelope purpurascens*), closely living with *O. vetula* within the infected area, did not show clinical signs. Some of these birds were sampled with cloacal and oral swabs, showing negative results to AIV.

It is also possible, that other avian species may have been infected as result of spillovers and not detected by the established active surveillance system. The complexity of the vegetation structure, may reduce the detection of ill birds. Also, the presence of opportunistic carnivores could have easily killed weak birds, including scavengers.

Although we do not know how the virus reached the reserve, we have established some possible routes of entrance. The first one is migratory birds in which many cases are the most common way of AIV entry into a bird population. The south bound neotropical migrants in El Zapotal, begins in July, reaching its peak during December and January [26]. During this migration there are no records of Anseriformes and Charadriiformes, the predominant reservoirs of AIV [26]. Also, during April 1-15, there was no accounts of mortalities, donations, confiscations or any other bird mobilizations in the ZooMAT.

The records of the visitors that attended the zoo during April 1-15, did not provide any useful information.

We do not discard that the interaction agent-bird could occur in ZooMAT in the food court area, where *O. vetula*, comes close to the people looking for food. The epizootic happened during the Holy Week period in which the number of visitors is very high. Because of this, it is possible that transmission might have occurred through fomites, mainly, shoes, clothes or contaminated material such as egg boxes that could have introduced the virus into the reserve [54]. Additionally, it is also possible that the holiday period incremented considerably the food consumption and sales in the restaurant, mainly eggs, originating product in the states under vaccination against H7 virus, recalling that the egg can be a source of infection [56, 58, 59, 60,].

During the breeding period, *O. vetula* may have had the chance to feed on cooked eggs and even egg shells from the garbage cans, becoming infected, considering garbage and containers, did not have an adequate manage [37]. Other possibility, could be the fights between males during the reproductive period, considering male-female ratio was 3 to 1. Once a bird is infected, the horizontal transmission of the virus is initiated.

Another form of transmission could have been related through foraging behavior of many birds. Some of them included an ample range of tropical fruits eaten by *O. vetula, T. grayi*. Although, considering the captivity of *Amazona albifrons*, the possibility of transmission could have happened due to human contamination. No hypothesis could be proved in this particular case. However, the results of the phylogenetic analysis inferred that the VIA H7N3 detected has an origin in the Mexican state of Puebla, second producer of egg for plate [60].

In 2004, a low pathogenic H5N2 influenza virus was identified in a psittacine bird for the first time in the United States under the Mexican lineage H5N2 viruses [20].

In addition, wild birds interact with poultry in a close vicinity. If these birds were AIV carriers, they contaminated the environment through oral and fecal excretions, and possibly carcasses [2, 7]. This environmental situation also increases the exposition and viral transmission risks to resident wild birds, known as “bridge birds” (BB). The BB can be infected by direct contact with feces or contaminated water, since the AI virus can survive in water during weeks or months [8, 57, 58, 59, 61]. Later, the BB will disseminate the virus by mechanical or biological means into the backyards or directly to poultry farms lacking of biosecurity (protections against synanthropic birds) [4, 8, 44, 58, 62].

This outbreak is significant because it shows the vulnerability of sites devoted to the protection and conservation of avian species, some of them under a risk category [28]. It is crucial to develop more complete analysis of the existing risks, and of the challenges to implement minimal biosecurity measures in these facilities. If the epizootic control had been unsuccessful, the viruses could have caused additional damage to other wild birds’ species as well as to the regional poultry industry with the subsequent loss of food production and jobs in one of the poorest states of Mexico.

Because of the susceptibility shown by *O. vetula*, its particular characteristics and wide distribution in the tropical and subtropical zones in Mexico and Central America, we recommend that this species, performs the role “as sentinel”. So, any event of mortality in this species, should be analyzed for Avian Influenza.

Finally, it is pivotal that veterinarians, wildlife managers and biologists notify to the veterinary authority, any suspicious activity of ill or event of mortality in wild birds, to be investigated [20, 25, 26].

## Acknowledgments

We wish to thank the Senasica Mexico, The Secretary of Environment and Natural History at the State of Chiapas, and the Regional Zoo Miguel Alvarez del Toro.

